# A hybridoma-derived monoclonal antibody with high homology to the aberrant myeloma light chain

**DOI:** 10.1101/2021.05.19.444780

**Authors:** Ghasidit Pornnoppadol, Boya Zhang, Alec A. Desai, Anthony Berardi, Henriette A. Remmer, Peter M. Tessier, Colin F. Greineder

## Abstract

The identification of antibody variable regions in the heavy (V_H_) and light (V_L_) chains from hybridomas is necessary for the production of recombinant, sequence-defined monoclonal antibodies (mAbs) and antibody derivatives. This process has received renewed attention in light of recent reports of hybridomas having unintended specificities due to the production of non-antigen specific heavy and/or light chains for the intended antigen. Here we report a surprising finding and potential pitfall in variable domain sequencing of an anti-human CD63 hybridoma. We amplified multiple V_L_ genes from the hybridoma cDNA, including the well-known aberrant Sp2/0 myeloma V_K_ and a unique, full-length VL. After finding that the unique V_L_ failed to yield a functional antibody, we discovered an additional full-length sequence with surprising similarity (~95% sequence identify) to the non-translated myeloma kappa chain but with a correction of its key frameshift mutation. Expression of the recombinant mAb confirmed that this highly homologous sequence is the antigen-specific light chain. Our results highlight the complexity of PCR-based cloning of antibody genes and strategies useful for identification of correct sequences.

## Introduction

Monoclonal antibodies (mAbs) are arguably the most important and widely used reagents in contemporary biomedical research and laboratory medicine(1, 2). They form not only the basis of numerous clinical tests, from lab based ELISAs to bedside lateral flow assays, but also commonly performed research techniques, including immunohistochemistry, western blotting, and flow cytometry. Unlike clinical antibodies, which are almost always produced as recombinant proteins in mammalian expression systems, biomedical research mAbs are typically prepared from culture of the original hybridoma cell lines(3). Since purification is typically accomplished via Fc-dependent affinity chromatography (i.e., Protein A or G) rather than methods which require affinity for the intended antigen, hybridoma-derived mAbs may be contaminated by other immunoglobulins secreted by these cell lines(4–6). The expression of heavy and light chains by myeloma fusion partners was a major concern for early hybridomas, which inevitably secreted antibodies with multiple paratopes and off-target specificities(7). The problem was mitigated by the creation and widespread adoption of Sp2/0, an immunoglobulin non-producing cell line(8), and related myelomas (e.g. NS-1, NS0, P3/X63Ag8.653, etc.), but hybridomas created using these fusion partners continue to express high levels of unproductive myeloma light chain mRNA(9, 10). In fact, multiple reports indicate the existence of additional heavy and/or light chain transcripts in many cell lines, some of which are capable of translation and production of full-length protein(11, 12). Prolonged culture, repeated passaging, and transfer between labs are likely to contribute to this problem, fueling speculation of widespread genetic and secretory variability in different preparations of supposedly “monoclonal” hybridomas(13, 14).

These suspicions have been confirmed using next generation sequencing (NGS), revealing a diversity of heavy and light chain transcripts in many hybridomas(14). While some investigators have called for NGS of all commercially available hybridomas and replacement with validated and standardized recombinant antibodies(15), the technology remains expensive enough to prohibit its widespread application. Instead, PCR-based cloning, in which degenerate primers are used to amplify heavy and light chain variable regions from hybridoma cDNA, remains the primary means of antibody gene sequencing for most laboratories(11, 16).

In our previous work, we reported a novel set of primers for PCR-based cloning (“SP primers”), which anneal to sequence encoding the signal peptide, just upstream of V_H_ and V_L_ coding regions(17). Unlike the more commonly cited “FR1 primers”, which are homologous to the first framework region, SP primers avoid the possibility of FR1 amino acid substitutions from incorporation of primer sequence(18, 19). The SP approach has another significant advantage – namely that the Sp2/0 myeloma light chain is typically amplified by just one SP primer, leaving the remaining PCR reactions available for amplification of antigen-specific sequences(17). We have used this approach to sequence more than 15 hybridomas with relatively few complications(20–23).

We report here an interesting case of PCR-based cloning using SP primers, in which identification of the antigen-specific light chain was complicated because it was significantly (~95%) homologous to the Sp2/0 myeloma kappa chain. The existence of this sequence is important to document because of to the ease with which it (or similar sequences) may be overlooked, as it was initially here. Likewise, methods aimed at eliminating the Sp2/0 myeloma sequence would likely destroy this cDNA and others like it(24, 25). Beyond cloning difficulties, the similarity to the myeloma light chain is intriguing because of the possibility that its origin might be a mutation of the myeloma sequence, rather than a splenic B cell. Ultimately, the work illustrates the complexity of PCR-based cloning of antibody genes and highlights a series of techniques capable of identifying antigen-specific sequences, even when initial efforts are confounding.

## Materials & Methods

### Cell lines

The mouse anti-human CD63 hybridoma, clone H5C6, developed by J.E. Hildreth at Johns Hopkins University(26), was obtained from the Developmental Studies Hybridoma Bank, created by the NICHD of the NIH and maintained at The University of Iowa, Department of Biology, Iowa City, IA 52242. CHO-K1 cells were obtained from ATCC (Manassas, VA). Both cell types were maintained in RPMI-1640 supplemented with 10% (v/v) fetal bovine serum (FBS), and 1X antibiotic-antimycotic (Thermo Fisher Scientific, Waltham, MA). For production of antibody, the hybridoma was gradually transitioned to Protein Free Hybridoma Medium II (PFHM-II, Thermo Fisher Scientific) and cultured for several days until the medium turned acidic. Supernatant was purified using Protein G Sepharose 4 Fast flow (GE Healthcare Life Sciences, Pittsburgh, PA) as per manufacturer protocol.

### Sequencing of H5C6 V_H_ and V_L_ cDNAs

Total cellular RNA was isolated from hybridoma cells using RNeasy kit (Qiagen, Valencia, CA). Reverse transcription was performed using High-Capacity cDNA Reverse Transcription Kits (Applied Biosystems, Foster City, CA). V_H_ and V_L_ cDNAs were amplified using the ProFlex PCR system (Thermo Fisher Scientific and a previously reported set of degenerate signal peptide (SP) region primers(17).

### N-terminal protein (Edman) sequencing

Purified antibody from H5C6 hybridoma supernatant (3 ug) was reduced in beta-mercaptoethanol containing sample buffer and run on SDS-PAGE. The gel was transferred to a PVDF membrane and stained in 0.1% Coomassie R-250 in 40% methanol and the band corresponding to the light chain was excised. N-terminal amino acid sequencing was performed using the Procise 494HT Edman sequencer (Applied Biosystems, Foster City, CA) with 610A data analysis module. Each amino acid was identified by comparison of its HPLC trace to a standard set.

### Assembly, production, purification, and characterization of recombinant antibodies

V_H_ and V_L_ cDNAs were cloned into a heavy and light chain vectors, both of which were in HEK293-6E expression vectors (pTT5, National Research Council Canada) with human IgG1 constant domains. A sortag (amino acid sequence LPETGG) was fused to the end of each heavy chain to enable site-specific modification with the sortase A enzyme(22). All recombinant antibodies were expressed via transient transfection with 7.5μg of each heavy and light chain plasmids (15μg total) in HEK293-6E suspension cells maintained with FreeStyle-17 media (supplemented with glutamine, kolliphor, and geneticin) using polyethylenimine transfection reagent. Supernatant was collected 6 days later and recombinant mAb was purified using protein A agarose (Thermo Fisher Scientific) according to the manufacturer’s protocol.

### Fluorescence labeling of hybridoma-derived and recombinant antibodies

H5C6 hybridoma-derived antibody was modified with AlexaFluor 647-NHS ester (Thermo Fisher Scientific) at a 1:5 (antibody: NHS-ester) ratio for 1 hour at room temperature in PBS. Modified antibody was purified from the reaction mixture using a Amicon 100kDa MWCO centrifugal filter (MilliporeSigma, Burlington, MA). Recombinant antibody, in contrast, was site-specifically modified using a previously reported technique for sortagged affinity ligands(22). Briefly, each recombinant mAb was incubated overnight at room temperature with 1μM A5 mutant sortase A enzyme(27), 1mM CaCl_2_, and a 5-fold excess of an azidolysine containing -GGG peptide. Azide-modified antibody was first purified from the sortase reaction mixture using an Amicon 100kDa MWCO centrifugal filter and then reacted with a 5-fold excess of DBCO-AlexaFluor647 (Click Chemistry Tools, Scottsdale, AZ). Fluorescently labeled antibody was purified using centrifugal filtration. Degree of labeling for both hybridoma-derived and recombinant mAbs was calculated using their absorbance at 280 and 650nm.

### Human CD63-GFP expressing cells

The CD63-pEGFP C2 plasmid was a gift from Paul Luzio (Addgene plasmid #62964)(28). The plasmid was transfected into CHO-K1 cells using Lipofectamine 2000 per manufacturer protocol. Cells were selected in media containing 1μg/mL of Geneticin (Thermo Fisher Scientific) and flow sorted on EGFP expression using a MoFlo Astrios cell sorter (Beckman Coulter, Brea, CA).

### Cell binding and flow cytometry

Cho-hCD63-EGFP cells and Cho-K1 control cells were trypsinized, counted, and resuspended in 10% serum containing media at a concentration of 10^6^ cells/mL. Cells were incubated for 1hr on ice with varying concentrations of fluorescently labeled hybridoma-derived or recombinant mAbs and then washed x3 in flow buffer (PBS + 3% fetal calf serum) prior to analysis on a ZE5 flow cytometer (BioRad, Hercules, CA). Equilibrium binding affinities were calculated via nonlinear regression (One site – total and non-specific binding) using Prism 7.0 software (GraphPad Software, San Diego, CA).

## Results

### Amplification of H5C6 variable region cDNAs

To amplify full-length V_H_ and V_L_ sequences from H5C6 hybridoma total cellular cDNA, we performed PCR with previously reported 5’ “SP primers” (homologous to heavy or light chain signal peptides)(17) and appropriate constant region primers (C_H1_ or C_L_). Figure 1 shows the PCR products generated using SP primers. One major band is seen for V_H_ (primer SP4), whereas two bands are present for V_L_ (primers SP2 and SP7).

**Figure 1.**
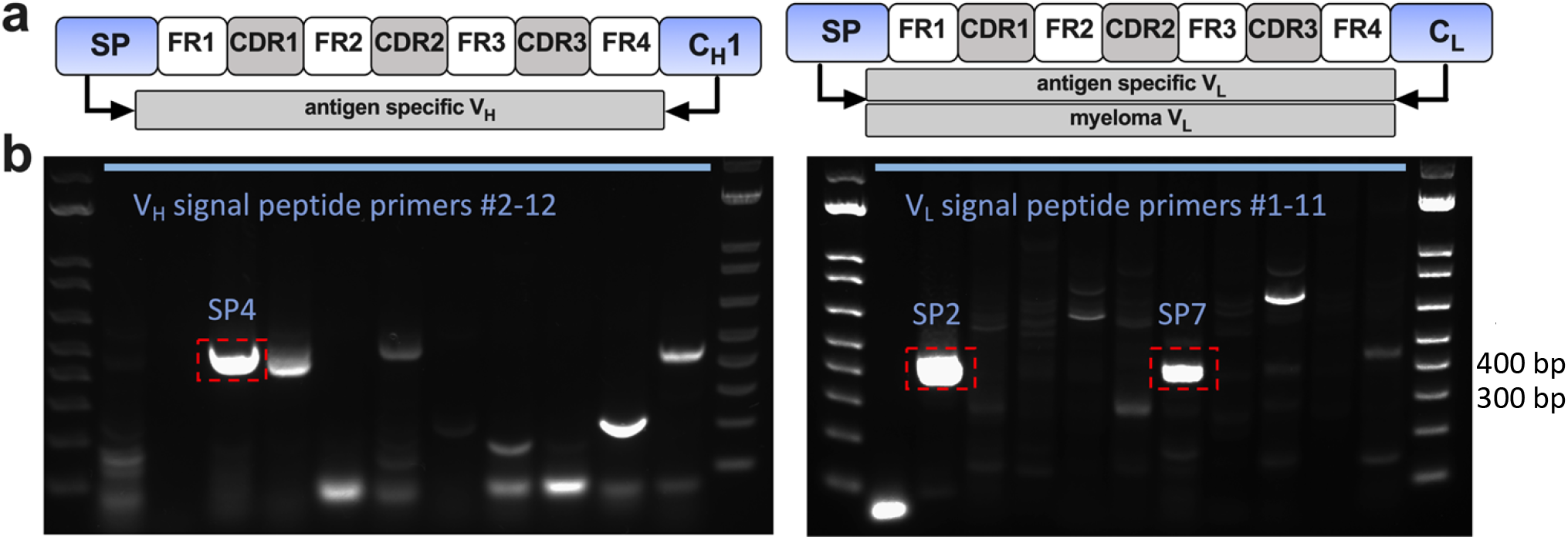
(a) Schematic of PCR amplification of VH and VL cDNAs by signal peptide (SP) primers. Right panel shows light chain PCR, which typically results in amplification of myeloma cDNA by some primers (primarily SP2) and antigen specific cDNA by other SP primers. (b) DNA gels. For the heavy chain (left panel), SP4 gave the major product, which was sequenced and revealed a full-length V_H_ cDNA. For the light chain (right panel), SP2 amplified the non-productive Sp2/0 myeloma kappa chain, whereas SP7 primer amplified a full-length V_L_ cDNA.

As with all prior Sp2/0-derived hybridomas investigated by our group, the SP2 primer amplified the aberrant myeloma kappa chain, a well-described sequence which does not produce secreted light chain due to the presence of a key frameshift mutation(10). While this transcript frequently complicates cloning using FR1 primers(11, 29–31), the results shown in Figure 1 are typical for SP primers, with amplification of the myeloma sequence limited to the SP2 (and, in rare cases, SP11) primer. Since other SP primers do not amplify the myeloma sequence, a properly sized amplicon in any of the other PCR reactions indicates a likely candidate sequence for the antigen-specific V_L_. Indeed, subcloning and sequencing of the SP4 and SP7 PCR products yielded single, full-length V_H_ and V_L_ cDNAs which were presumed to be hCD63-specific sequences.

### Human CD63 expressing cells and binding of recombinant mAbs

To evaluate binding to cell-surface hCD63, we generated CHO cells that stably express hCD63-EGFP fusion protein. We validated these using H5C6 hybridoma-derived mAb, purified from cell supernatant via Protein G affinity chromatography. Fluorescently modified antibody bound hCD63-expressing cells, but not wild type controls (Figure 2).

**Figure 2.**
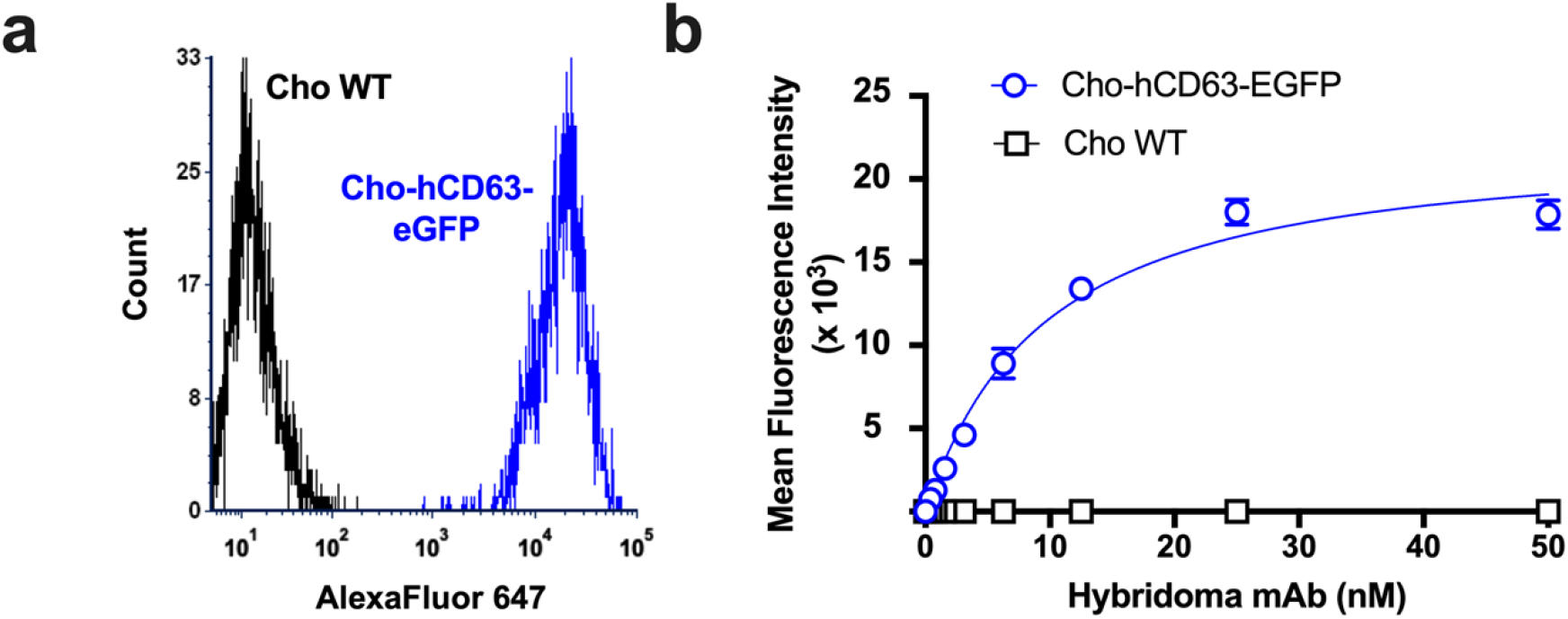
(a) Flow cytometry histogram in far red channel showing staining of Cho-hCD63-eGFP cells, but no Cho WT cells, with 25nM AF647-modified H5C6 hybridoma-derived mAb. (b) Binding curve derived from flow cytometry data (n=3).

We next generated recombinant mAb using the SP7 V_L_ and SP4 V_H_ sequences identified above (Figure 3a). SP7/SP4 mAb was purified from HEK293 cell supernatant and found to be the correct size and >95% purity on SDS-PAGE and size exclusion HPLC (Figure 3b, c). Recombinant mAb was modified for flow cytometry using an amine-reactive fluorophore, but it showed no binding to hCD63-expressing cells. While this strongly suggested the incorrect specificity, we considered the unlikely possibility that fluorescent labeling might have eliminated binding affinity by modification of one or more critical lysine side chains. To exclude this, we took advantage of the C-terminal sortag incorporated into the recombinant mAb construct, and site-specifically labeled the antibody using sortase(22, 32). Once again, no specific binding was found to hCD63-positive cells, as compared to wild-type control cells (Figure 3d, e).

**Figure 3.**
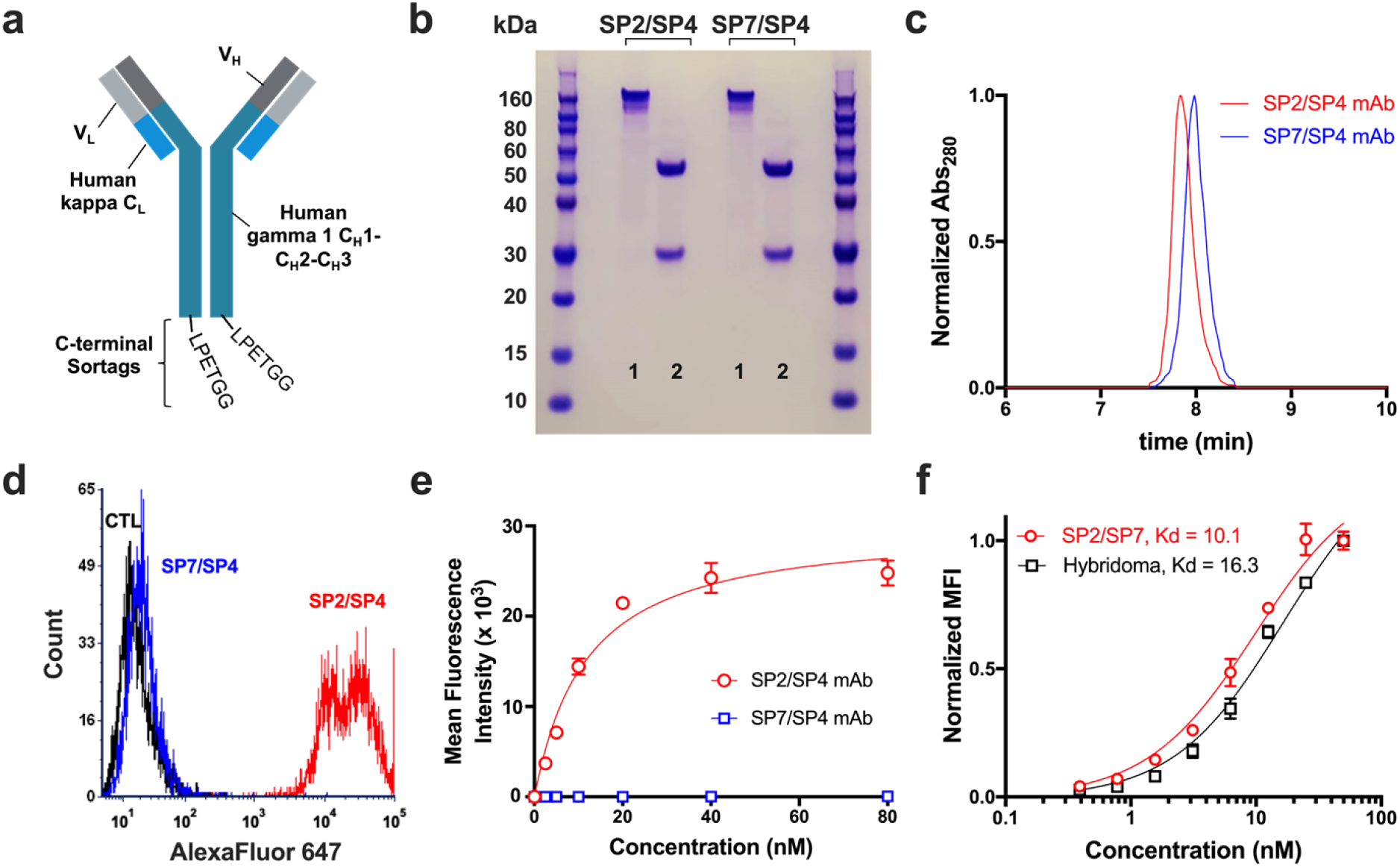
Characterization of recombinant mAbs. (a) Schematic of recombinant mAb constructs. VL sequences were cloned in frame with human kappa CL, while VH was cloned in frame with human CH1-CH3. The sortag sequence, LPETGG, was appended at the C-terminus, immediately after the CH3 domain. (b) SDS-PAGE of recombinant mAbs, 1 = non-boiled, non-reduced, 2 = boiled, reduced. (c) Size exclusion HPLC of SP2/SP4 and SP7/SP4 mAbs, showing a single peak on the A280 detector at the expected size for immunoglobulin (7.9 min). (d) Flow cytometry histogram showing binding of 20nM SP2/SP4 recombinant mAb, but not 20nM SP7/SP4 mAb, to Cho-hCD63-eGFP cells. (e) Flow-based binding curve no Cho-hCD63-eGFP cells (neither mAb showed any binding to Cho WT cells). (f) The affinity of SP2/SP4 mAb was similar to that of H5C6 hybridoma-derived monoclonal antibody (n=3).

### N-terminal protein sequencing and identification of an additional full-length V_L_ cDNA

Based on these data, we reasoned that another full length V_H_ and/or V_L_ sequence must be present in the H5C6 total cellular cDNA. To determine the N-terminal amino acid sequence of each chain and enable design of specific 5’ primers, we performed Edman sequencing on the H5C6 hybridoma-derived antibody preparation. The results for the light chain, shown in Figure 4, were unambiguous and showed that first 10 residues exactly matched those of the myeloma light chain (Figure 4A). This suggested the existence of a full-length, productive V_L_ cDNA with significant homology to the aberrant transcript, at least over the first 30 base pairs of the coding sequence. In fact, we hypothesized that the two cDNAs might have homology in the region of the signal peptide as well and that primer SP2 may have amplified them both. We subcloned the SP2 PCR product and, this time, sequenced a much larger number of clones. While many matched the Sp2/0 myeloma light chain, a few contained a full-length sequence with correction of the frameshift mutation. The new cDNA had a very high level of homology to the myeloma light chain, with 94% and 82% nucleic acid and amino acid sequence identity, respectively (Figure 4B).

**Figure 4.**
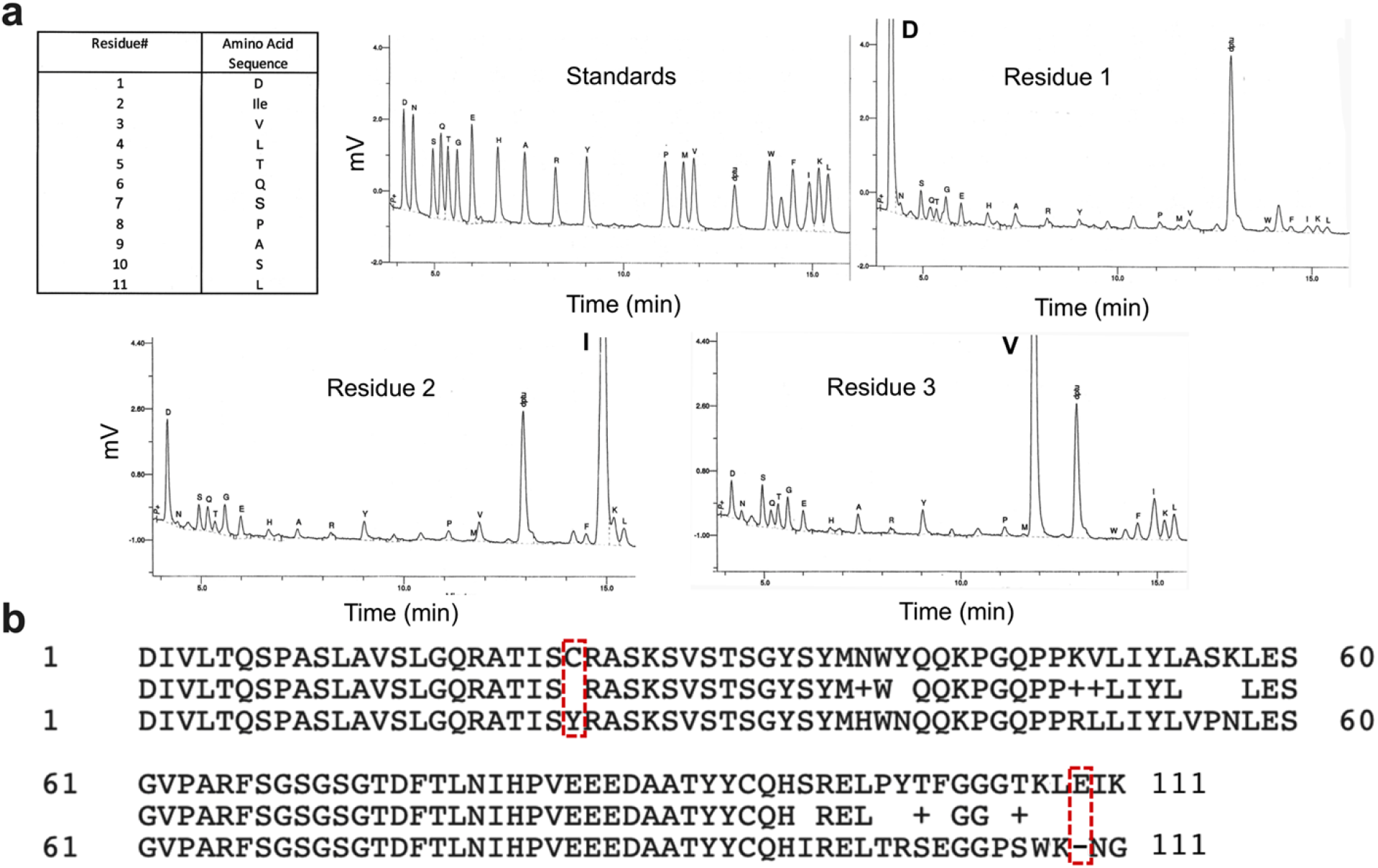
Identification of frameshift-corrected analog of Sp2/0 V_L_ cDNA. (a) Results of Edman sequencing of H5C6 hybridoma-derived mAb – summary, standards, and first three residues are shown. (b) Comparison of amino acid sequences of full-length SP2 V_L_ cDNA and aberrant Sp2/0 myeloma sequence. Both the frameshift mutation and the conserved cysteine (C23Y) mutation are corrected in the full-length gene product.

### Synthesis of recombinant mAb and confirmation of binding to hCD63

The newly identified SP2 V_L_ cDNA was incorporated into the recombinant mAb construct with SP4 V_H_. After confirmation of size and purity (Figure 3B), SP4/SP2 mAb was site-specifically labeled using sortase A and evaluated by flow cytometry for binding to hCD63-expressing cells. As shown in Figure 3B, the recombinant antibody showed high-affinity, saturable binding to Cho-hCD63 cells, but not wild-type controls. The affinity was compared to that of the hybridoma-derived mAb and found to be similar but slightly higher (K_D_ = 10.1 nM vs. 16.3 nM, Figure 3C).

## Discussion

This report represents a continuation of our ongoing efforts to sequence hybridomas with potential utility in affinity-based drug targeting and, in the process, to help refine and improve rapid, PCR-based cloning of antibody variable chain sequences(17, 21, 22). While some have advocated for next-generation sequencing (NGS) to replace traditional techniques, we believe that the substantial cost associated with this technique makes PCR-based methods a more viable option for most laboratories and justifies continued focus on this approach.

In our previous report, we detailed the use of signal peptide (“SP”) primers as a means of quickly and efficiently differentiating antigen-specific and myeloma-derived V_L_ sequences(17). Implicit in this approach is the assumption that these sequences do not share significant homology. Our experience with the H5C6 hybridoma is an important reminder that this assumption may not always be valid. It is worth noting that this pitfall is not unique to PCR-based hybridoma sequencing, as the high level of homology observed in this case likely would have confounded any technique. Even NGS, which would have presumably identified both sequences, would be unlikely to classify the full-length SP2 V_L_ as a candidate for antigen-binding. Rather, the sequence would likely be flagged as either a sequencing artifact or a frameshift-corrected version of the Sp2/0 sequence, perhaps resulting from prolonged hybridoma culture.

Given the significant challenge that this pitfall posed, the current results are also a demonstration of how a careful and systematic approach can result in successful identification of variable chain sequences, even when initial results are confusing. Several key steps can be identified as being of particular importance to our end result. First, our efforts benefitted from a transient expression system capable of rapid production of candidate mAbs and appropriate reagents for testing of antigen binding. The latter included a positive and negative cell line, hybridoma-derived mAb for comparison, and a site-specific modification technique to exclude potential artifacts related to fluorescent labeling(22). Together, these resources led to prompt recognition that the SP7/SP4 mAb was incorrect, while other assays – e.g., testing mAbs on tissue slices – might have delayed or obscured this conclusion. A second key feature of our approach was the use of SP primers, which typically produce a smaller number of V_L_ reactions with products of the correct size (presumably reflecting less overlap in the range of sequences capable of amplification by each primer). This reduces the burden of subcloning and screening multiple clones per fragment. In the case of H5C6, it pointed the finger clearly at the SP2 reaction, once the SP7 sequence had been excluded. If not for our prior experience with SP2, we likely would have screened a large number of SP2 clones at this point and successfully identified the full-length V_L_. Instead, it took another technique -- protein sequencing, which has been useful in working out other challenging hybridomas(20, 23). Ultimately, we believe that NGS will simplify this process considerably, but a systematic approach, high quality reagents, and an awareness of potential pitfalls will remain critical to successful identification of variable chain sequences.

Apart from these pragmatic observations, the current work should add to the growing chorus of scientific voices advocating for replacement of hybridoma-derived preparations with sequence, defined recombinant mAbs(5, 15). As an affinity-based drug targeting laboratory, these reagents are essential to nearly every aspect of our work, and clearly, they are not the immutable cell lines some consider them to be(14). Indeed, one of the most interesting questions raised by the current results is the origin of the hCD63-specific light chain. The original report of the hybridoma describes fusion of “splenic B cells from immunized mice with the P3×653.Ag8 myeloma”(26). While this name likely refers to the non-producing myeloma, P3/X63Ag8.653, it is also possible that Hildreth and colleagues inadvertently used P3/X63.Ag8, the fusion partner from Kohler and Milstein’s original publication, which secretes a fully functional kappa light chain(1). Still, this would not explain the presence of the non-translated Sp2/0 transcript in the H5C6 clone, which instead suggests a subsequent recombination or mutation that corrected the aberrant light chain. Moreover, none of this clarifies what may have happened with the light chain from the original splenic B cell. These considerations do underscore, however, the complexity and inherent instability of these tetraploid cells and the importance of their gradual replacement with sequence-validated, recombinant cell lines.

## Acknowledgements

This work was supported by the National Institute of Health grants K08 HL130430 (CFG), RF1AG059723 (PMT) and R35GM136300 (PMT), and the Albert M. Mattocks Chair (PMT).

## Authorship

G.P. and C.F.G. conceived the study. G.P. and A.D. designed and synthesized recombinant mAbs. C.F.G. and B.Z. made and purified antibody from the hybridoma and performed flow studies. C.F.G. and A.B. made and characterized the hCD63 expressing Cho cells. H.R. performed Edman sequencing. G.P., P.M.T., and C.F.G. wrote and edited the manuscript.

